# Cotranslational formation of disulfides guides folding of the SARS COV-2 receptor binding domain

**DOI:** 10.1101/2022.11.10.516025

**Authors:** Amir Bitran, Kibum Park, Eugene Serebryany, Eugene I. Shakhnovich

**Author notes:** These authors contributed equally to this work (Amir Bitran, Kibum Park).

## Abstract

Many secreted proteins contain multiple disulfide bonds. How disulfide formation is coupled to protein folding in the cell remains poorly understood at the molecular level. Here, we combine experiment and simulation to address this question as it pertains to the SARS-CoV-2 receptor binding domain (RBD). We show that, whereas RBD can refold reversibly when its disulfides are intact, their disruption causes misfolding into a nonnative molten-globule state that is highly prone to aggregation and disulfide scrambling. Thus, non-equilibrium mechanisms are needed to ensure disulfides form prior to folding in vivo. Our simulations suggest that co-translational folding may accomplish this, as native disulfide pairs are predicted to form with high probability at intermediate lengths, ultimately committing the RBD to its metastable native state and circumventing nonnative intermediates. This detailed molecular picture of the RBD folding landscape may shed light on SARS-CoV-2 pathology and molecular constraints governing SARS-CoV-2 evolution.

## 1 Introduction

In recent decades, a growing body of work has cast doubt on the generality of Anfinsen’s dogma of protein folding, which states that a protein’s amino acid sequence contains all the information needed to ensure folding to the native state [1]. Instead, myriad examples have emerged of proteins that cannot reliably refold from a denatured state on physiological timescales, instead populating highly stable nonnative intermediates that may be prone to aggregation [2–9]. In extreme cases, these intermediates may even correspond to global minima on the free energy landscape, whereas the native state represents a high-energy state–an energetic feature that may give proteins dynamic flexibility in response to stimuli or environmental changes [10–13]. For such proteins, which cannot fold under thermodynamic control, two major questions arise: 1.) How do cellular mechanisms ensure the native state is attained under kinetic control, and 2.) What factors ensure that the native state persists over biological timescales despite the existence of alternative stable states?

A relevant feature of many proteins, pertaining to the above questions, is the presence of disulfide bonds[14]. These covalent linkages between cysteine residues can confer as much as 5-6 kcal/mol of stability per bond by reducing the entropy of the denatured state, and thus significantly increase the population of the native state. [15–17]. At what stage during protein folding do these highly stabilizing disulfide bonds typically form? For certain proteins, the correct disulfides can spontaneously form following dilution from a denatured, reduced state into a suitable redox buffer [1, 18–20]. In these cases, the non-covalent free energy landscape readily guides the protein to a native-like structure in which the cysteines are suitably positioned for spontaneous oxidation and disulfide formation. In other cases, the underlying folding landscape may favor the native state, but kinetic intermediates along the folding pathway may promote temporary nonnative disulfide formation, or even random disulfide scrambling [21, 22]. These nonnative bonds can then isomerize into the native disulfides, either unassisted or with the aid of enzymes such as protein disulfide isomerase [23–25]. But in a certain subset of proteins, the native disulfides cannot be reformed at all starting from a denatured, reduced state, as the underlying energy landscape strongly favors alternative folds that are structurally incompatible with these disulfides[22, 26, 27]. Such proteins must necessarily rely on cellular mechanisms to ensure the correct native disulfides are present prior to folding.

In the cell, many proteins can begin forming their disulfide bonds co-translationally as they are being secreted into the endoplasmic reticulum (ER). The ER is a highly oxidizing environment and contains multiple enzymes such as protein disulfide isomerase (PDI) and Ero1 that catalyze disulfide formation in substrates [28–30]. Numerous studies have shown that co-translational folding in the ER can significantly enhance the efficiency of native folding and disulfide formation while reducing the formation of nonnative intermediates relative to post-translational oxidative refolding [23, 27, 31–34]. This improvement may stem from the vectorial nature of synthesis, which can allow native structural units to form sequentially and thus mitigate misfolding that arises post-translationally [35–41]. However, the specific molecular details underlying these processes remain poorly understood in general.

In this work, we address these questions as they apply to the receptor binding domain (RBD) of the spike protein in SARS-CoV-2. This virus, which is behind the Covid-19 pandemic, has killed millions of people worldwide to date and continues to wreak havoc on lives and economies across the globe [42]. Thus there is a pressing need for novel antiviral therapies that can broadly target novel, highly contagious variants as they emerge. The SARS-CoV-2 RBD, which comprises amino acids 333 through 527 of the S1 subunit of the spike protein, contains multiple disulfide bonds which have previously been shown to be crucial for the ensuring domain’s stability and functional binding to the human ACE2 receptor [43, 44]—an interaction that directly leads to viral-host fusion [45, 46]. However, the mechanism through which RBD disulfide formation is coupled to its folding has not been investigated.

Using fluorescence assays, we show here that the RBD can only be refolded reversibly from denaturant if its native disulfide bonds are present prior to refolding. In contrast, oxidative refolding from a denatured and fully reduced state leads to misfolding into a nonnative molten-globule like state that is highly prone to aggregation. This implies that the native state is kinetically trapped by its disulfides, and that the intrinsic free energy landscape of the reduced RBD favors alternative nonnative structures. Using a combination of gel-based assays, we then show that, upon oxidation, the nonnative structures form fewer disulfide bonds than the native state. Atomistic simulations recapitulate these findings and generate predictions of specific disulfides that are most likely to be absent in this state. These simulations additionally predict that, at intermediate lengths during RBD synthesis, native cysteine pairs frequently come into contact while nonnative pairs typically remain far apart suggesting that co-translational oxidation in the endoplasmic reticulum may help commit the RBD to its metastable native state. Together, these results paint a detailed molecular picture of how disulfide formation is coupled to folding in a biomedically crucial model protein. This study may in turn further our understanding of SARS-CoV-2 pathology and molecular evolution and inspire novel antiviral therapies that interfere with RBD folding.

## 2 Results

### RBD can only refold reversibly if native disulfides are present prior to folding

To begin, we investigated whether the RBD can refold reversibly from a denatured state, depending on whether or not the domain’s native disulfide bonds are disrupted during denaturation (Fig 1a). To this end, we chemically denature the RBD in 5 M guanidine hydrochloride (GdnHCl) in the presence or absence of 1 mM TCEP as a disulfide reducing agent, then refold the protein in oxidizing buffer containing 5 mM oxidized glutathione. The refolded protein’s intrinsic fluorescence spectrum was then compared to the native protein’s to assess if refolding was successful.

**Fig. 1.**
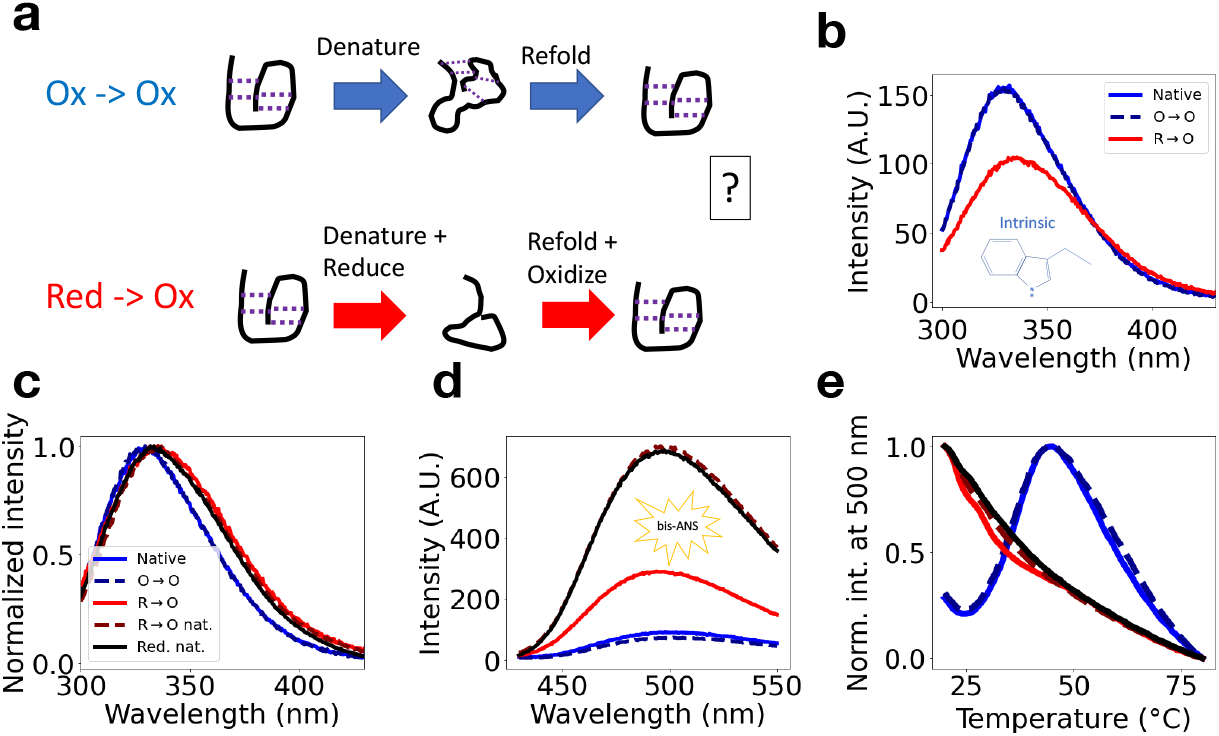
RBD can only refold reversibly in presence of disulfides. **a**. Schematic of Ox →Ox and Red→ Ox protocols. RBD is denatured in 5 M GdnHCl in the presence (Red →Ox) or absence (Ox →Ox) of 1 mM TCEP to reduce disulfide bonds. Refolding is then initiated via 10-fold dilution into oxidizing buffer containing 5 mM oxidized glutathione. For details, see main text and Methods. **b**. Intrinsic fluorescence spectra (excitation wavelength of 280 nm) of 1.2 *µM* refolded RBD prepared via the Ox →Ox (labeled O →O)and Red →Ox (labeled R →O) protocols, as well as 1.2 *µM* native protein in 0.5M GdnHCl to match final denaturant concentration after refolding. Each spectrum represents an average of two independent replicates. **c**. Same spectra as in b. normalized to maximum emission value to emphasize peak shift, alongisde normalized spectra of 1.2 *µM* Red→Ox native sample in which reduction was performed under native conditions (labeled R→O nat.) and sample reduced under native conditions that was not re-oxidized (labeled Red. nat.), both in 0.5M GdnHCl. All unnormalized spectra are shown in Supplementary Fig. 1c. **d**. bis-ANS fluorescence spectra (excitation wavelength of 385 nm) for all samples in panel c. The bis-ANS concentration is 8 *µM*. The colors and linestyles match those in panel c. **e**. bis-ANS fluorescence at 500 nm (385 nm excitation wavelength) as a function of temperature for samples prepared as in fig. b-d. The colors and linestyles match those in panel c.

We observe that, if denaturation is performed in the absence of reducing agent—a protocol we refer to as Ox → Ox to reflect that both denaturation and refolding occur under oxidizing conditions—then the refolded protein’s intrinsic spectrum perfectly overlaps with that of the native state (Fig 1b). Thus, refolding is reversible provided the native disulfides are maintained intact during denaturation. But if the denatured protein is reduced with TCEP then re-oxidized during refolding–a protocol referred to as Red→ Ox– then the refolded protein’s fluorescence spectrum shows a pronounced red-shift and decrease in intensity relative to the native state (Fig 1b-c). This finding, which is robust to the composition of the glutathione buffer used during refolding (Supplementary Fig. 1) suggests that the refolded, re-oxidized protein adopts a nonnative conformation in which the RBD tryptophan residues (two per subunit) are more exposed to solvent than in the native state. As a second structural probe for refolding, we incubated all constructs with bis-ANS–a dye that fluoresces upon binding hydrophobic cavities–and measured the resulting fluorescence. Strikingly, we observe that the Red → Ox construct shows significantly higher bis-ANS fluorescence at room temperature than the native and Ox → Ox constructs, suggesting that the former may exhibit molten-globule like character whereas the latter constructs are well-folded (Fig. 1d). To further confirm this, we measured the dependence of bis-ANS fluorescence on temperature for each construct (Fig. 1e). The native and Ox→Ox melting curves show non-monotonic fluorescence with temperature that peaks at 50°*C*–an indicator of a well folded protein in which hydrophobic residues are compact at room temperature but become looser and more exposed at intermediate temperatures prior to complete denaturation. These two curves overlap, further indicating that the Ox→Ox leads to native refolding. In stark contrast, the Red→Ox construct shows monotonically decreasing fluorescence, with half-maximal intensity occurring around 37°*C*. This further points towards nonnative molten-globule like character in Red→Ox, and hints that this construct may be prone to high-temperature aggregation resulting in loss of fluorescence–similar behavior has previously been observed in destabilizing mutants of the enzyme dihydrofolate reductase [47].

These results indicate that RBD can only attain its correct structure if its native disulfides are present prior to folding, whereas disulfide reduction during denaturation followed by oxidative refolding leads to misfolding. One mechanism that may explain this misfolding is if, early during refolding, the chain becomes locked into kinetically accessible, but high energy states via semirandom disulfide formation—a behavior previously observed in proteins such as hirudin [21, 28] and RNAse-A [48]. To assess this possibility, we determined whether similar nonnative misfolding occurs if the protein is reduced in the absence of denaturing agent–under these conditions the protein should only populate states that are low in free energy under native conditions. Indeed, this preparation produces a nearly identical intrinsic fluorescence red-shift to that observed in the Red → Ox construct (Fig. 1c) as well as increased bis-ANS fluorescence (Fig. 1d) and a monotonically decreasing melting curve (Fig. 1e). These results, which are observed whether or not this reduction under native conditions is followed by re-oxidation (samples referred to Red → Ox native and Reduced native, respectively), rule out the possibilty that that these nonnative states correspond to trapped high-energy intermediates, and indicate that folding is thermodynamically irreversible upon disruption of disulfides Interestingly, the *magnitudes* of the bis-ANS and intrinsic fluorescence in these Red→Ox native and Reduced constructs differ from those in Red→Ox (Fig. 1d and Supplementary Fig. 1c), suggesting that these protocols may lead to slightly different non-native structures–in the case of reduced native, such a difference is expected given that the protein was not re-oxidized.

### RBD oxidative refolding produces a stable nonnative intermediate that is highly aggregation prone

Our observation that the refolded, re-oxidized RBD is more hydrophobic than the native state and shows molten-globule like behavior hint that this species may be prone to aggregation. To test this, we ran our Red → Ox species, as well as native RBD on a size-exclusion chromatography (SEC) column (Fig. 2a) We find that, whereas the native RBD elutes primarily as a monomer as expected (MW~ 35 kDa accounting for N-linked glycans), the Red→ Ox sample largely elutes near the column void volume suggesting it is forming large oligomers whose molecular weight exceeds ~ 100 kDa (the exclusion limit for our Superdex 75 column.) Notably, centrifuging this Red → Ox sample prior to loading does not produce any visible precipitate, suggesting these oligomers are soluble.We observe a similar void peak if protein is prepared via the Red → Ox Native preparation (Supplementary Fig. 2), and this sample shows fluorescence upon incubation with Thioflavin T (ThT)—an indicator of amyloid-like aggregation—to a significantly greater extent than the native protein (Supplementary Fig. 3a-b). Interestingly, we note that our standard Red → Ox preparation in which RBD is denatured during reduction does not lead to appreciable ThT fluorescence, perhaps because residual denaturant during refolding affects the aggregates’ structure such that they are no longer amyloid-like (Supplementary Fig. 3c). Together, these findings indicate that the oxidative refolding causes RBD to populate a molten-globule intermediate that is highly prone to forming soluble aggregates.

**Fig. 2.**
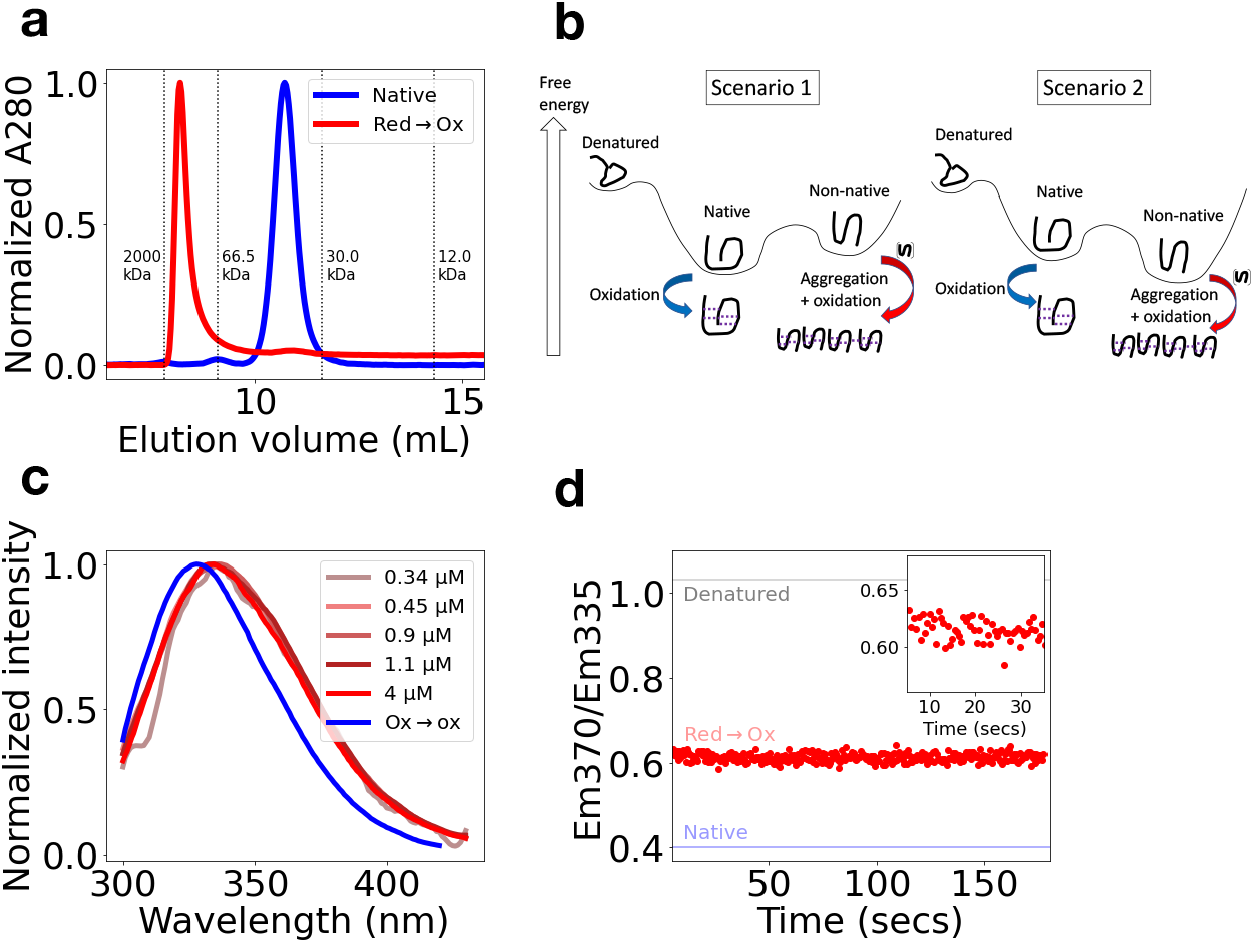
Reduced RBD folding landscape intrinsically favors a highly aggregation-prone nonnative intermediate. **a**. Size-exclusion chromatography UV absorbance traces, normalized to maximum value, for native RBD and refolded RBD per Red →Ox protcol, defined as in Fig 1. Dashed vertical lines indicate elution positions for standards with indicated molecular weights. **b**. Two possible scenarios that may explain observed misfolding and aggregation. Solid curves with three minima represent energy landscapes for RBD monomers in the absence of disulfides. For details, see main text. **c**. Normalized intrinsic fluorescence spectra for refolded Red Ox →RBD samples at final protein concentrations indicated, alonside Ox →Ox sample (1.7 *µM*). **d**. Ratio of intrinsic fluorescence intensity (solid red dots) at 370 nm to intensity at 335 nm as a function of time following manual-mix refolding (5 second dead time) averaged over two traces. The final protein concentration is 0.5 *µ*M. The solid blue and gray lines depict the expected final ratio for RBD in the native and denatured + reduced state, respectively. The inset shows the refolding kinetics zoomed in on early times.

We next sought to relate this observed aggregation to the folding landscape of RBD monomers. Our findings thus far are consistent with two possible scenarios (Fig. 2b):

1. The native state represents the free energy minimum in the reduced RBD’s folding landscape, and thus most molecules spontaneously adopt a native-like state upon refolding. However, transient structural excursions into a high-energy intermediate, which occur before the native state can become locked in place via disulfide formation, lead to aggregation. Disulfide formation then further locks the monomers in this nonnative state.
2. The aggregation-prone monomeric intermediate (likely an ensemble of rapidly interconverting molten-globule structures) represents the minimum free energy state in the *reduced RBD’s* folding landscape, and thus most of the population spontaneously adopts it upon refolding then proceeds to aggregate, followed by disulfide formation as per scenario 1. A third scenario, that the aggregation results from the monomers adopting an energetically unfavorable yet highly kinetically accessible state upon refolding, can be ruled out by our previous finding that reduction under native conditions leads to similar molten-globule like behavior (Fig. 1c-e) and substantial aggregation (Supplementary Fig. 2). Notably, in both of the above models, the fully oxidized, native RBD’s free energy is lower than that of any state on the reduced energy landscape and thus, native oxidation can effectively be treated as an absorbing boundary–however aggregation of the nonnative RBD represents a second absorbing boundary that prevents the native state from being reached in our experiments.

To distinguish between scenarios 1 and 2, we can run refolding experiments under a wide range of RBD concentrations to effectively modulate the aggregation timescale relative to the oxidation timescale, which is constant at a fixed redox buffer composition. If the aggregation process is dependent on transient fluctuations away from the native state per scenario 1, then reducing the protein concentration (and thus aggregation rate) should decrease the protein population that ultimately becomes locked in aggregates prior to oxidation, thus increasing the proportion of correctly-folded protein and yielding a more native-like fluorescence spectrum. However this is not what we observe, as the final (normalized) fluorescence spectra are essentially indistinguishable from each other over an order-of-magnitude range of protein concentrations (0.34 *µ*M to 4 *µ*M, Fig. 2c). This striking insensitivity to concentration points in favor of scenario 2.

As a further test of scenario 2, we directly monitored the kinetics of refolding by measuring the ratio of fluorescence at two wavelengths—370 nm and 335 nm—as a function of time following manual mixing of denatured protein with refolding buffer. This fluorescence ratio is expected to acquire a value of 0.4 in the native state, whereas in the nonnative state it is closer to 0.6 and in the denatured, reduced state it is around 1. Strikingly, the vast majority of the refolding amplitude occurs within the dead time of manual mixing (5 secs) with only minimal kinetics afterward (Fig. 2d). By 10 seconds, the signal has stagnated at the expected nonnative value of 0.6. These kinetics, which are qualitatively similar if an alternative wavelength pair is monitored (320 and 370 nm, see Supplementary Fig. 4a), are much too rapid to correspond to aggregation at the protein concentration used (0.5 *µ*M) This is because the rate of aggregation cannot be any faster than the diffusion-limited association rate, which for most proteins is in the range of 10^5^ *−* 10^6^ M^*−*1^*s*^*−*1^. Protein pairs with evolved complimentary charged interfaces can associate faster than this owing to electrostatic steering forces, but in the case of RBD, the monomers have a net +7 positive charge at physiological pH. Thus we expect that the 105 *−* 10^6^ M^*−*1^*s*^*−*1^ range represents an upper bound on the association rate for RBD, which at submicromolar protein concentrations, translates to a 20-200 second timescale. The observation that the kinetics are nearly complete by 5 seconds and stagant after 10 seconds strongly suggests that this fluorescence ratio is not reporting on aggregation at all, but rather on nonnnative tertiary structure within the monomers. Thus, this nonnative monomer structure is substantially populated even prior to aggregation, strongly pointing towards scenario 2.

In additional support of scenario 2, we observe similarly rapid kinetics, and a similar nonnative final ratio if both denturation and refolding are performed under *reducing* conditions (Supplementary Fig. 4c). This, along with our finding in Fig. 1c-e that reduction is sufficient to produce a molten-globule like intermediate, indicates that the folding landscape of the reduced RBD favors nonnative monomeric states. Furthermore upon reducing native RBD, we observe a substantial separation between the timescales for intrinsic fluorescence change, which is rate-limited by disulfide reduction and is largely complete within an hour, and aggregation as monitored by ThT fluorescence, whose rate is highly-concentration dependent and generally exhibits a multi-hour lag phase (Supplementary Fig. 4d-3). This is consistent with our expectation from scenario 2 whereby relatively rapid and spontaneous misfolding of monomers should precede slower aggregation.

### Nonnative RBD structure shows incomplete disulfide bond formation

Given that RBD can correctly refold in the presence of its native disulfides (Ox → Ox) whereas disruption and re-formation of these disulfides leads to misfolding (Red → Ox), we expect that the misfolded, re-oxidized structure should exhibit a nonnative pattern of disulfide bonding. One possibility is that the refolded monomers contain fewer disulfide bonds than the native state, whose disulfide bonding pattern is shown in Fig. 3a. If so, then the additional free cysteines in the misfolded state may mediate disulfide cross-links between aggregated monomer subunits. To test for such cross-links, we ran both our Ox→ Ox and Red → Ox on a denaturing SDS-PAGE gel either with or without reducing agent (5 mM TCEP) added prior to loading the respective sample (Fig. 3b), as well as iodoacetamide to alkylate any free cysteines and prevent disulfide scrambling. Interestingly, we observe that, whereas the Ox → Ox sample runs as a monomer independent of TCEP, the Red → Ox sample runs as a ladder of mixed oligomeric species which disappears when TCEP is added–very similar results are observed using Red → Ox native samples (Supplementary Fig. 5a). This TCEP-dependence indicates that the observed higher-order oligomers are indeed disulfide-linked. We thus conclude that the misfolded monomers must contain at *minimum* one fewer disulfide than the native state, which only has one free cysteine and thus cannot form cross-linked species larger than dimers. We note that, in addition to these intermolecular disulfides, the RBD oligomers must also be held together by non-covalent interactions that are disrupted by SDS denaturation–otherwise we would similarly observe a ladder while running the sample through SEC under native conditions rather than a single peak at the void volume (Fig. 2a). We expect that under oxidizing conditions, these noncovalent interactions between subunits form prior to nonnative disulfide crosslinks, although further work is needed to characterize the precise mechanism of aggregation.

To further characterize the difference in disulfide bonding between the samples, we incubated both in the presence of PEG-maleimide–an alkylating reagent that conjugates free cysteines to a bulky PEG group, while leaving disulfide-paired cysteines intact. By running both on a gel and observing mobility shift, we find that the Red → Ox sample shows 3 4 additional PEGylation bands than Ox → Ox, suggesting that the former contain up to two fewer disulfides than the latter (Supplementary Fig. 5b-d). However, the relatively low PEGylation efficiency under our conditions, which we attribute to aggregation and disulfide cross linking reducing the number of free cysteines, as well as a potentially inhomogenous starting population render it difficult to obtain a clearly quantifiable difference between the constructs with this assay.

### Atomistic simulation model for nonnative state aligns with experimental observations

Next, we investigated whether atomistic simulations can generate a detailed structural model for the RBD nonnative states that is consistent with experimental observations. To this end, we made use of the DBFOLD method–an all-atom enhanced-sampling Monte-Carlo simulation platform and analysis pipeline that is uniquely capable of sampling both native and nonnative folding intermediates not accessible to conventional simulations on reasonable computational timescales [49, 50]. We start off by running simulations of RBD with or without distance constraints between native cysteine pairs, which mimic the effect of disulfide bonds (see Methods). Although our simulation force field is not currently equipped to model RBD’s N-linked glycans, we note that these glycans have been primarily linked to spike protein expression and cellular processing [51], and their absence does not appear to alter RBD structure, ACE2-binding, or immunogenicity [52]; likewise we do not expect their omission from simulations to drastically affect our findings.

Following the completion of simulation runs, we asked whether the equilibrium folding landscapes with and without disulfide constraints favor different structures. To describe the folding landscape, we generate a contact map of the native RBD structure, and subdivide the map into islands of proximal native contacts that are expected to form cooperatively, known as substructures (Fig. 4a). By computing the free energy of states associated with different permutations of formed/broken substructures, known as *topological configurations* [35, 49, 53], we can determine the energetics associated with forming and breaking different structural units at equilibrium. Notably, we observe that whereas the folding landscape in the presence of disulfide constraints favors the fully folded state, the unconstrained landscape instead favors a nonnative configuration in which the interfaces between four beta strands are disrupted, namely the strands comprising residues 3-5, 27-30, 58-60, and 190-192 (Fig. 4b). Thus, these simulations agree with the experimental finding that, in the absence of disulfides, the non-covalent energy landscape favors alternative non-native states over the native state. To assess whether the disruption of these beta sheets is due to absence of disulfide constraints, we can directly monitor the probability that different cysteine pairs make contact with each other in these unconstrained simulations (Fig. 4e). Consistent with our gel-based cross-linking and PEGylation assays (Fig. 3b and Supplementary Fig. 5), we find that the nonnative state typically contains 1-2 fewer cysteine pairs than the native state, and indeed, the two most frequently missing pairs in the absence of constraints (Cys 3-28 and 58-192) occur at the interfaces of the disrupted beta sheets (Table 1). Thus, per simulation, the RBD folding landscape favors conformations that bring together some, but not all native cysteine pairs. Interestingly, our simulations predict that the the unconstrained folding landscape brings together a *nonnative* cysteine pair, 28-58, with substantially higher probability (30%) than either of the native pairs involving these two cysteines, namely 3-28 and 58-192 (which occur with probabilities 2% and 15%, respectively). This suggests that the nonnative state may be prone to nonnative disulfide scrambling–a prediction that can be directly tested in future work.

**Fig. 3.**
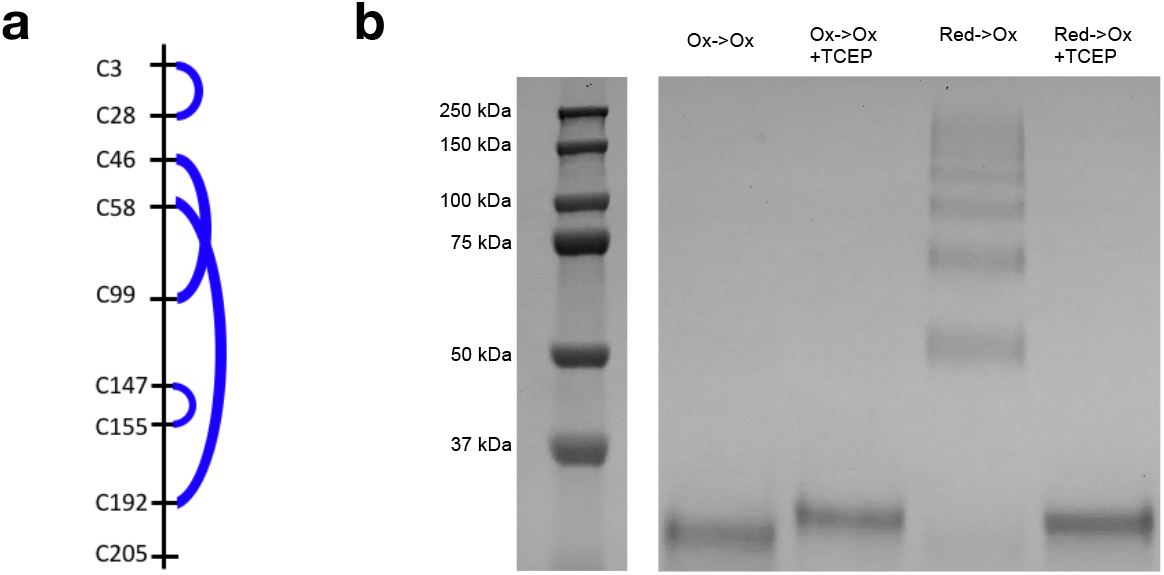
RBD nonnative state shows incomplete disulfide formation and is prone to forming cross-linked aggregates. **a**. Schematic of cysteines and disulfides (blue curves) in RBD. In our numbering scheme, residue 1 of the RBD corresponds to residue 333 of the full spike protein. **b**. SDS-PAGE gel, stained with coomassie G-250 dye, loaded with Ox →Ox and Red →Ox refolded samples with or without 10 mM TCEP added prior to loading, as indicated above respective lanes. Inset on left shows labeled molecular weight standards.

**Fig. 4.**
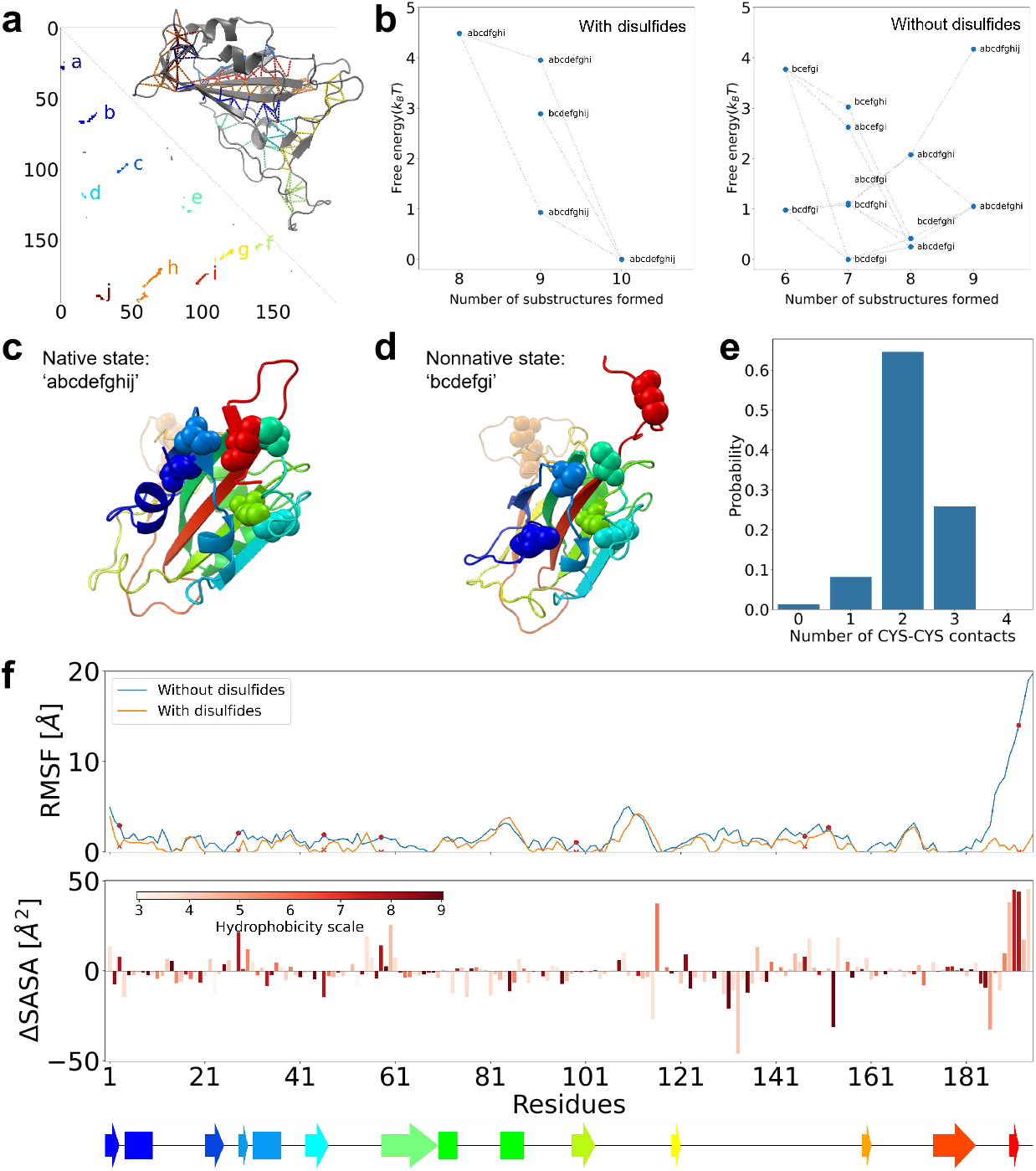
Atomistic simulations explain experimentally-observed structural properties of misfolded RBD. **a**. Native contact map for RBD, alongside native structure. Contacts corresponding to the substructures *a* to *j* are indicated with different colors and labeled on structure as dashed lines. **b**. Free energies of distinct *topological configurations*, defined as the collection of snapshots containing some subset of folded native substructures (defined as in a). Each dot represents one topological configuration, as labeled, with its x-position corresponding to its respective number of folded substructures and y position corresponding to its respective free energy relative to the lowest energy state. **c**. Example snapshot of fully-folded structure (configuration *abcdefghij*) from the simulation with constraints. **d**. Example snapshot of configuration *bcdefgi*, the lowest energy state from simulations without disulfide constraints.Native disulfide bonds between CYS46(cyan) and CYS99(light green) and CYS147-155(orange) are still present while CYS3, 28, 56, and 192 do not form disulfide bonds. **e**. Probability distribution of number of cysteine-cysteine contacts among snapshots from the simulation without disulfide constraints **f**. Root mean square fluctuation relative to native state (top plot) of each residue in simulations with disulfide constrains (orange) and without constrains (blue line, cysteine positions indicated with red markers), and difference in solvent-accessible surface area (SASA) between the simulations as a function of residue (bottom plot). Positive ΔSASA indicates the residue is more exposed in the simulation without constraints. The color of each bar represents the hydrophobicity of the residue, quantified by the Miyazawa-Jernigan scale [75]. Secondary structure of each residue is indicated below the plots (arrows = beta strands, rectangles = helices, colored as per Fig. 4c-d)

**Table 1.**
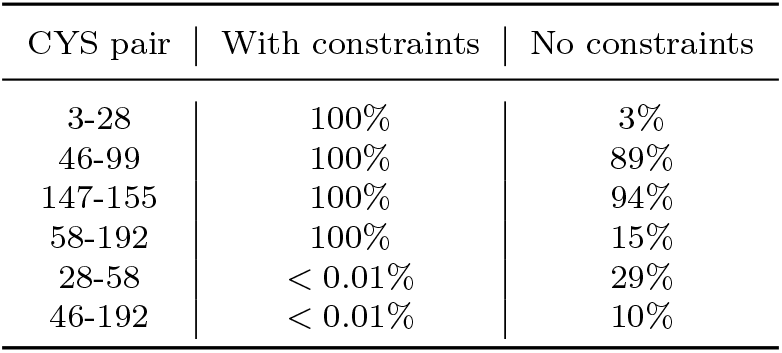
Probabilities that indicated cysteine pairs come together within a beta-carbon distance of 5*Å* in full-length RBD simulations with and without disulfide constraints.

| CYS pair | With constraints | No constraints |
| --- | --- | --- |
| 3-28 | 100% | 3% |
| 46-99 | 100% | 89% |
| 147-155 | 100% | 94% |
| 58-192 | 100% | 15% |
| 28-58 | < 0.01% | 29% |
| 46-192 | < 0.01% | 10% |

Our atomistic simulations further generate specific structural predictions that may explain why the nonnative state shows molten-globule like behavior and high aggregation propensity. Namely, we observe that the majority of residues are markedly more dynamic (i.e. show higher root-mean-squared fluctuation with respect to the native state) in the absence of constraints–this discrepancy is especially pronounced at the N and C-terminal beta strands but some degree of difference is observed throughout, even in segments without cysteines. (Fig. 4f, top). Furthermore, certain hydrophobic patches are more exposed in the unconstrained simulation, most notably the disrupted C-terminal beta strand, but also to some extent the disrupted N-terminal strand as well as core segments around residues 25-30, 55-61, 108-110, and 116-121. This loosened, more exposed protein core may explain the enhanced bis-ANS fluorescence in the nonnative state, and the highly dynamic N and-C-terminal beta strands may help mediate intermolecular aggregation (Fig. 2a), ThT fluorescence (Supplementary Fig. 3, often an indicator of amyloidgenic aggregation between beta strands), and intermolecular cross linking via the free cysteines (Fig. 3). We note however that, although it shows partial molten-globule like character, the nonnative state does not behave as a classical molten globule as it contains numerous ordered, well-packed regions in common with the native state. In line with this, we observe that not all hydrophobic residues are more exposed in the nonnative state–in particular, two long stretches containing multiple hydrophobic residues between AAs 62-115 and AAs 119-153 are generally *less exposed* in this alternative state. Interestingly, this latter stretch largely coincides with the disordered ACE2 binding interface, hinting that these functionally important segments may introduce energetic frustration into the protein. In addition to exposed hydrophobics, this binding interface also contains multiple positively charged residues that are closely spaced in the native state, potentially introducing further energetic strain to the native state.

### Simulation model predicts that co-translational folding improves RBD disulfide-formation and folding efficiencies

Together, our experimental and simulation results indicate that the fully-synthesized RBD cannot be refolded from a denatured reduced state as the underlying folding landscape favors nonnative, molten-globule like states with incomplete/incorrect disulfides and a significant aggregation propensity. Given that many proteins can begin folding co-translationally, we wondered whether disulfide formation during RBD secretion into the ER may help ensure correct disulfides form, ultimately locking the protein into the native state and improving folding efficiency. To investigate this, we used our all-atom DBFOLD algorithm to simulate RBD constructs truncated at various chain lengths shortly after the synthesis of successive cysteine residues. For each length we assessed the probability that native and nonnative cysteine pairs come into contact (within a beta-carbon distance cutoff of 5 Angstroms), and thus can be oxidized into a disulfide.

From these simulations (Fig. 5a), we find that most native cysteine pairs show an appreciable probability of coming into contact at intermediate lengths. Perhaps the most salient feature of these profiles is that the 3-28 pairing probability varies non-monotonically with length, peaking around length 100 (~20% probability) before declining to less than 5% at the full length (195 residues). This is indicates that, whereas the full-length RBD is highly prone to adopting misfolded states that keep this cysteine pair apart, the energy landscape at intermediate lengths is more likely to bring this native pair together. In contrast, *nonnative* cysteine pairs rarely come together at intermediate lengths.

**Fig. 5.**
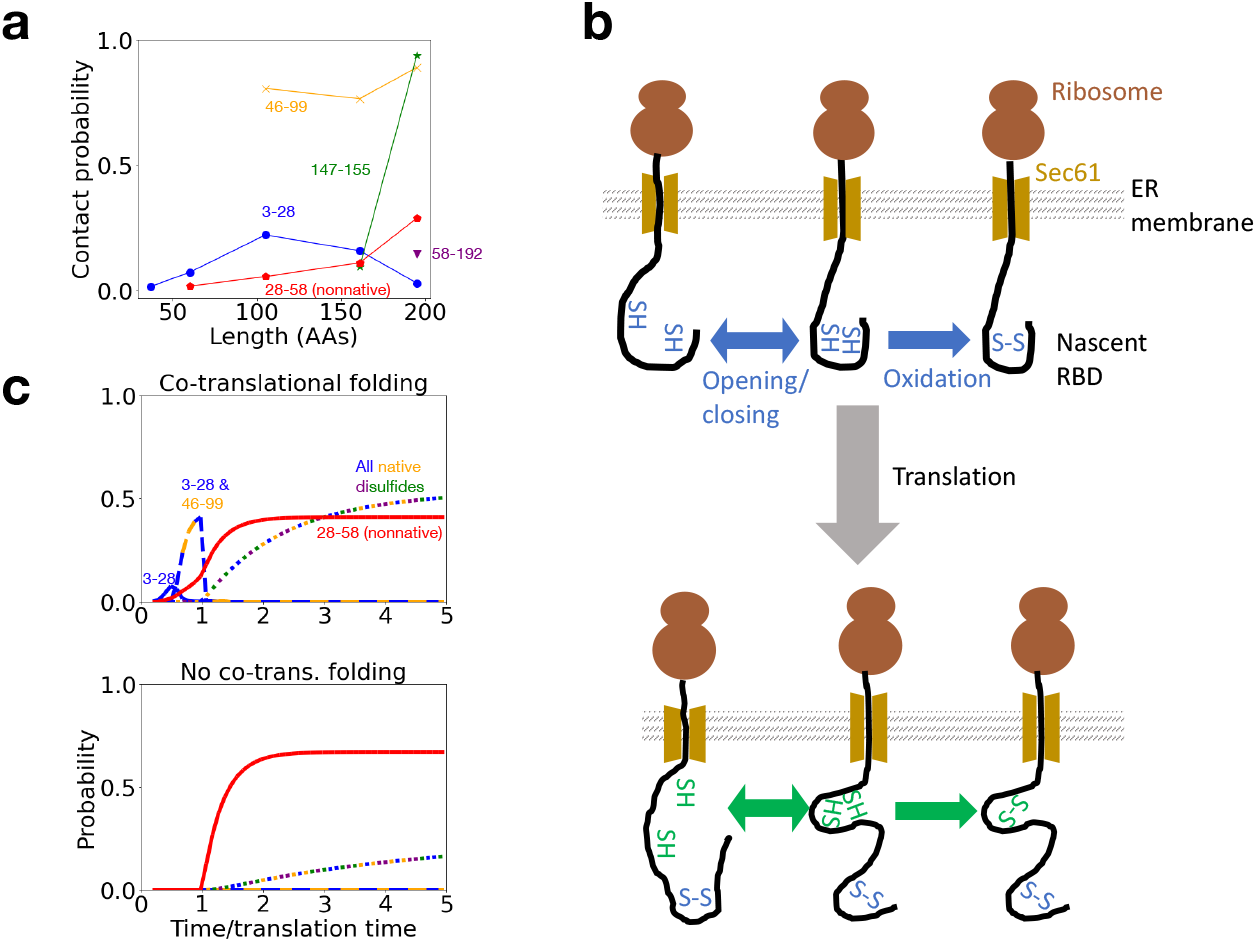
Atomistic simulations predict that co-translational folding significantly improves native RBD formation rate and yield. **a**. Probability, as a function of translation length, that cysteine pairs (as indicated in legend) make contact within 5 Angstroms of each other as predicted by atomistic simulation. **b**. Schematic of kinetic model, detailed in the main text and Methods sections. In our kinetic model, opening and closing transitions that unite or spearate cysteine pairs (double arrows) are assumed to occur on much faster timescales than oxidation(single arrows pointing right) (per equation (1)). At a given length, any permutation of new disulfides may form, but only on such formation sequence at the first two lengths is depicted here. **c**. Solutions to kinetic model showing probability, as a function of time, of forming native species containing various permutations of the four native disulfides (indicated above respective curves) as well as trapped species containing the 3-28 nonnative disulfide (red line), assuming either that folding begins co-translationally per the kinetic model in b (top panel), or does not begin until synthesis is complete (bottom panel). The x-axis depicts time normalized by the time needed to translate the full RBD–we note that synthesis must be complete before the final dislufide involving C-terminal residue 192 can form. In this panel, the unknown intrinsic oxidation to translation rate ratio, *kox/ktrans* is set to 5.

The most likely such pair, namely 28-58, rarely occurs prior to length ~ 150, but this probability increases at later chain lengths, and by 195 residues (the full length), this nonnative pair probability exceeds that of the native 3-28 pair. (other nonnative pairs are almost never observed at all–see Table 1)

A key parameter that will affect the efficiency of co-translational disulfide formation is the ratio between the characteristic timescale for translation of the domain (40 100 seconds, assuming a translation rate of 2-5 AAs/second [54])and the characteristic timescale of disulfide formation by PDI–which has been shown to be capable of catalyzing disulfide formation co-translationally [28–30]–provided a cysteine pair is in contact. To quantify the effect of this parameter, and the advantage to co-translational folding more generally, we constructed a kinetic model of co-translational disulfide formation (Fig. 5b, Methods). The key assumptions underlying this model are:

1. At any given length *L*, the rate at which a given disulfide *d* forms is independent of whether other disulfides are present. This is justified by control simulations which show that, at a given length, the presence of all disulfides that could have formed prior to that length does not significantly affect probability of additional cysteines coming into contact (Supplementary Fig. 8).
2. A disulfide bond can only form if the respective cysteines first come into proximity of each other–here we use a distance cutoff of 5 Angstroms. After such a cysteine-cysteine contact is formed, the pair is oxidized at a rate *k*_*ox*_ that is assumed to be insensitive to the identity of the cysteines. In the cell, PDI is expected to catalyze this oxidation step.
3. Disulfide formation is assumed to be an irreversible process. Although in reality disulfide shuffling can occur, this process is typically quite slow compared to translation timescales and is thus neglected here.
4. The rates of structural fluctuation that bring cysteines into contact are assumed to be much faster than the oxidation rate *k*_*ox*_. This assumption is in line with previously measured co-translational folding rates on the millisecond timescale [55] compared to minute timescales for PDI catalytic activity [24]. Thus, combining this assumption with the first one, we can express the effective rate of formation for disulfide *d* as:

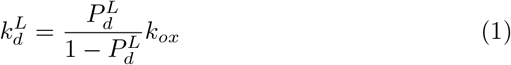

Where 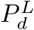 is the probability that the cysteine pair corresponding to *d* makes contact at length *L*, as per Fig. 5a, and the above ratio thus represents the equilibrium constant for these cysteines coming together. We can then write down a master equation at length *L* which governs the time evolution for the probability of occupying states with different permutations of formed disulfides:

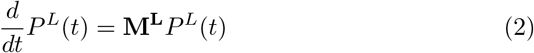

where *M*_*L*_, the transition matrix at length *L*, incorporates the (irreversible) disulfide-formation transitions with rates obtains as described above. We note that in addition to proximity, the rate of disulfide formation is also affected by the cysteine pair’s surface accessibility [30]–however we observe that, among snapshots where a given cysteine pair is formed, the pairs’ total SASA shows minimal dependence on chain length, and the SASA values the two competing cysteine pairs that are most likely to determine folding outcome (3-28 and the nonnative 28-58) fall within a relatively narrow (∼2-fold) range (Supplementary Fig. 9). Thus, for simplicity, we neglect the effect of accessibility on catalytic rates.

For a given ratio of the characteristic oxidation rate to the translation rate for the full domain (*k*_*ox*_*/k*_*trans*_, a free parameter), we can solve for the probability, as a function of time, that species with different numbers of cysteine pairs have formed, either assuming that co-translational disulfide formation occurs, or that disulfides cannot form until synthesis is complete. To begin, we set *k*_*ox*_*/k*_*trans*_ = 5. Assuming a cysteine pairing probability of ∼20%, this would yield an effective disulfide formation rate equal to the RBD translation rate, which is reasonable given that both processes should occur on minute timescales. Under this condition we observe that co-translational folding leads to a roughly ten-fold speedup in the initial rate of native product formation following the synthesis of the final cysteine pair. Furthermore, the final native yield is roughly two-fold higher when co-translational folding is permitted, and the amount of nonnative product (containing the 28-58 disulfide) reduced. This is because, with co-translational folding, the nascent chain can take advantage of intermediate chain lengths at which the native 3-28 pair is significantly more likely to form than the competing 28-58 pair–a trend that is reversed at the full length where the 3-28 pairing probability is extremely low and nonnative disulfides are more likely (Fig. 5a). In Supplementary Fig. 10a, we vary the oxidation-to-translation ratio over a reasonable range and show that this benefit to co-translational folding is robust to the exact value, although the benefit is naturally increased at higher oxidation rates. We note that this trend is somewhat more sensitive to another unknown parameter–the simulation temperature (which, in the context of our coarse-grained potential, cannot be precisely translated to real temperature)–but some degree of benefit is predicted at a wide range of temperatures below the melting transition (Supplementary Fig. 10b)

## 3 Discussion

By combining in vitro experiments with atomistic simulation, our work generates a unique, detailed molecular picture of the SARS-CoV-2 co-and-post translational oxidative folding landscapes. These findings advance our understanding of how protein folding is coupled to disulfide formation more broadly. In some proteins, the formation of native secondary and tertiary structure precedes the disulfide bonds, and the underlying folding landscape strongly favors native over nonnative cysteine pairings–such proteins fold via a so-called structured precursor mechanism [1, 18–20, 23, 28, 30]. But this mechanism is at odds with our finding that RBD can only be refolded if its native disulfides are maintained intact during denaturation, whereas disruption of these disulfides leads to misfolding and nonnative disulfide formation. Thus, the fully-synthesized RBD more closely resembles proteins that fold via alternative mechanism, known as the quasi-stochastic model, in which the folding landscape intrinsically favors nonnative states with significant disulfide heterogeneity [22, 24–27]. Our simulation model agrees with our experimental finding that the RBD nonnative state contains fewer disulfides than the native state, and additionally predicts the specific native cysteine pairs–namely 3-28 and 58-192–that are most likely to be disrupted, as well as the possibility of the nonnative 28-58 disulfide. These predictions can be tested using mass-spectrometry based assays. Our experimental and simulation results further indicate that these nonnative states are molten-globule like and highly prone to aggregation– similar aggregation-prone molten globule states have been observed in certain destabilizing mutants of the enzyme DHFR [47, 56], as well as other systems. This aggregation tendency further suggests that the presence of other spike protein domains may further exacerbate the folding deficiencies observed here for isolated RBD, given that interdomain misfolding and nonnative interdomain disulfide formation are oft-observed risks [23, 41, 57, 58]. These effects are not included in our kinetic model and may even further heighten the benefit of co-translational folding.

For proteins with a significant nonnative disulfide propensity, co-translational folding during secretion into the ER may help improve the speed and efficiency of native disulfide formation and folding. Our atomistic simulations suggest that the co-translational oxidative folding of RBD proceeds via a hybrid between the folding-precursor and quasi-stochastic models. On the one hand, our simulations predict that at intermediate translation lengths, the folding landscape significantly favors native over nonnative disulfide formation. This is in stark contrast to other systems in which co-translational intermediates show significant propensity towards nonnative disulfides [23, 27, 34]. But despite this bias towards native disulfides, our simulations predict that the co-translational intermediates are not fully ordered, and native secondary/tertiary structure does not fully form until after synthesis, provided the disulfides have begun forming co-translationally (Supplementary Fig. 7). In this sense, the RBD’s co-translational mechanism more closely resembles the quasi-stochastic model, in which completion of folding typically follows disulfide formation. These structural predictions can be tested *in vivo* using pulse-chase assays [31–33], or using *in vitro* translation systems alongside pulse-labeling techniques such as hydrogen-deuterium exchange [59]. Furthermore, our simulations predict that disrupting co-translational oxidation of the early-forming 3-28 disulfide bond in particular (e.g. via circular permutation) will drastically decrease the native folding efficiency and increase the population of misfolded proteins, some fraction of which will contain the scrambled 28-58 disulfide.

Our results here may also shed further light on previous findings that reduction of RBD’s disulfides leads to altered secondary and tertiary structure, reduced stability, and impaired ACE2 binding [43, 44]. Consistent with this, it is known that expressing RBD in E. coli leads to a disulfide-lacking species that is highly aggregation-prone and must be refolded from inclusion bodies [52, 60]. One possible mechanism to explain these observations is that, in the absence of correct disulfides, RBD is more prone to structural fluctuations that expose hydrophobic segments, leading to aggregation (Scenario 1 in Fig. 2), as obsered in disulfides-containing proteins such as hen lysozyme and SOD1 [61, 62]. But our work points to a more striking mechanism—namely that the native structure represents a metastable state that is disfavored thermodynamically in the absence of disulfides (Scenario 2 in Fig. 2). Thus, RBD cannot efficiently fold under standard conditions, and is expected to benefit significantly from non-equilibrium mechanisms that promote folding under kinetic control. Although this work emphasizes the role of co-translational folding, we note that a previous study showed that RBD can successfully be expressed in *E. coli* –an expression system with limited disulfide formation capacity–and subsequently oxidized in suitable buffer, provided the RBD is fused to a solubility-enhancing peptide (SEP) tag [52]. However, this finding does not contradict our results here, as the key reason why the native state cannot be reached in our experiments is due to irreversible aggregation of the nonnative state. If this aggregation is suppresed, as may be achieved by a SEP tag, then native, oxidized state will correspond to the global free energy minimum and eventually be reached. However, the nonnative state corresponds to the free energy minimum on the *reduced* RBD’s energy surface–thus upon refolding, this state is rapidly adopted initially and subsequently partially oxidized to form two out of four native disulfides in most molecules, and two native and one nonnative disulfide in a fraction of molecules (Fig. 2b scenario 2 and Fig. 4). Sufficient suppression of aggregation could provide time for this nonnative state to undergo structural fluctuations to high energy states that bring together the remaining cysteine pairs, which subsequently get oxidized, and for any nonnative pairs to isomerize, ultimately yielding the global minimum native state. Similar suppression of aggregation may also be achieved via dropwise refolding of resolubilized RBD inclusion bodies–another method shown to produce folded, soluble protein following E. coli expression [60]. But in the crowded cellular environment, aggregation and interdomain misfolding represent significant hazards, suggesting that co-translational folding is a more likely mechanism to ensure efficient native folding.

A key outstanding question left by this work is, what factors contribute to the stability gap between the metastable RBD native state and the low-energy nonnative state we observe, and could these factors paradoxically be linked to RBD’s function? One pertinent observation from our atomistic simulations is that disordered segments in RBD tend to exhibit higher flexibility in the nonnative state than in the native state, indicating that conformational entropy may be important in stabilizing the former (Fig. 4F). Yet these disordered regions also form a key region of the ACE2 binding interface [45, 63–67] and are conserved across evolution [67], strongly pointing towards a functional role—perhaps this disorder increases the domain’s avidity towards ACE2 by allowing for multivalent interactions [68]. These disordered regions are further enriched with positively charged residues that allow the RBD to form strong ionic interactions with the negatively charged binding interface [45]. But this cluster of like charges may introduce energetic frustration to the native state, and indeed, we observe that nonnative structures tend to be more expanded with these charges further apart. Additional studies are needed to confirm these physical tradeoffs between folding and function, and determine how they may shape the evolution of SARS-CoV-2 variants with different ACE2 binding affinities. We further note that, in addition to our finding that the RBD structure represents a metastable state, it is known that the pre-fusion conformation of the spike protein as a whole is likewise kinetically trapped [69]. This metsastability allows the spike to undergo a functionally crucial conformational change into the lower-energy post-fusion state. However in the case of the RBD, we do not anticipate our observed metastability to be functionally relevant as the lower energy state is highly aggregation prone.

Finally, our work may lay the groundwork for future therapeutic directions against SARS-CoV-2. Recent studies have suggested that abnormal blood clotting, caused by fibrin aggregation, may be linked to various Covid-19 longterm sequelae [70, 71]. Notably, such fibrin aggregates have been shown to contain spike protein molecules, hinting that misfolded or aggregated spike protein can nucleate fibrin aggregation [72]. It would be interesting to examine whether variability in redox metabolism across patients could modulate the folding and disulfide-formation efficiency in spike protein, and thus affect subsequent aggregation and clinical outcome. This knowledge can also be used to improve efficiency and reduce side effects of vaccines that use the RBD as an antigenic agent. In the alternative scenario that the misfolded RBD state is clinically benign, our study can inspire the design of folding inhibitor drugs that selectively stabilize nonnative structures and promote RBD misfolding, thus inhibiting virus-host membrane fusion and replication.

## 4 Methods

### Preparation of refolded samples

The receptor binding domain from SARS-CoV-2 S1 protein] (amino acids 319-541), expressed in Hek293 cells and purified to ≥ 95% purity, was purchased from Thermo Fisher (RP87704). The lyophilized protein in PBS buffer was reconstituted to a concentration of 0.5-1 mg/ml (∼ 20*µM*) in MilliQ water and the purity was confirmed by SDS-PAGE. The protein then denatured in in 5 M guanidine hydrochloride, pH 7.4 supplemented with either 1 mM of tris(2-carboxyethyl)phosphine hydrochloride (TCEP, Research Products International) to produce the Red →Ox samples, or an equivalent volume of MillliQ water to produce the Ox→ Ox samples. The protein concenentration was then adjusted, as necessary, either via dilution in respective denaturing buffer, or concentration with Amicon Ultra 0.5 mL centrifugal filters. To achieve maximal protein concentration (in Fig. 2c), the protein was lyophilized protein was directly reconstituted directly in 5M GdnHCl buffer. After a minimum of 6 hours incubation under denaturing conditions, the protein was refolded via 10-fold dilution in either 5 mM oxidized glutathione (Millipore 3542) or 3.3 mM oxidized glutathione + 1.7 mM reduced glutathione dissolved in PBS buffer, pH 7.4 for at least 6 hours to overnight prior to measurement. For samples that were to be loaded on a gel (as in Fig. 3), the denaturation step was performed with 8M urea instead of GdnHCl–we show in Fig. S1 that both denaturants produce very similar final spectra following refolding.

To produce the native control, Red → Ox, native, and reduced samples (Fig. 1), the protein was prepared identically as above, except that during the first incubation step, the denaturant was replaced with an equivalent volume of PBS buffer, pH 7.4 supplemented with 5 mM TCEP (in Red → Ox native, and Reduced samples) or an equivalent amount of water (in native control). The native control and Red→Ox samples were then diluted in 5 mM glutathione buffer spiked with GdnHCl for a final concentration of 0.5 M to match the final concentration in refolding measurements, whereas the reduced sample was diluted in PBS buffer spiked with TCEP and GdnHCl for final concentrations of 5 mM and 0.5 M, respectively. As before, both steps were performed for a minimum of 6 hours.

### Fluorescence measurements

For steady state intrinsic fluorescence measurements, a minimum of 70 *µ*L of protein sample was transferred into a 1 cm Helma fluorescence cuvette. The cuvette was then loaded into a Varian Cary Eclipse Fluoresecence Spectrophotometer, and the sample was excited at 280 nm with a 5 nm excitation slit width and a 10 nm emission slit width at a scanning speed of 150 nm/min, averaging time of 0.2 seconds, and data collection interval of 0.5 nm. A blank measurement of respective buffer + protein replaced by an equivalent volume of PBS was also acquired. Samples were then recovered, and 4,4’-Dianilino-1,1’-Binaphthyl-5,5’-Disulfonic Acid, Dipotassium Salt (bis-ANS) was added to a final concentration of 8 *µ*M. The samples were incubated for 10-15 minutes to allow the dye to bind the protein, and the resulting bis-ANS fluorescence was measured using an excitation wavelength of 385 nm. All other measurement parameters were the same as those used for intrinsic fluorescence, and a blank, bis-ANS containing sample was similarly prepared and measured. All data processing, including blank correction and normalization to maximum, was carried out using custom Python scripts.

For thermal denaturation assays, protein and blank samples were prepared with 8 *µ*M bis-ANS as above. The bis-ANS fluorescence (*λ*_*ex*._ = 385 nm, *λ*_*em*._ = 500 nm) was then measured over a temperature range from 20 °*C* to 80 °*C* using a scanning speed of 0.5°*C/*min and 1 second averaging time. A blank sample was also measured, although it showed negligible change in fluorescence with temperature compared to the samples. For each sample, fluorescence at all temperatures then divded by the maximum value observed to produce normalized melting curves (as in Fig. 1d).

The kinetics measurements were initialized with an empty cuvette in the cell holder, with an excitation wavelength of 280 nm and emission wavelengths of 335 nm and 370 nm (1 ms integration time, excitaiton slit 5 nm, emission slit 10 nm). At 1 second, the denatured protein was mixed with 90 *µ*L of refolding buffer (5 mM oxidized glutathione in PBS, pH 7.4), and immediately transferred to the cuvette. The dead time, accounting for the time needed to transfer refolded protein to cuvette and initial mixing artifacts, is 5 seconds.

### Size exclusion chromatography

Proteins were separated on a Superdex 75 Increase 10/300 GL column (Cytiva) that was pre-equilibrated with PBS buffer, pH 7.4. Samples were prepared at 9 *µ*M and centrifuged at 15,000 g for 15 minutes to remove any insoluble aggregates. The supernatant (200 *µ*L) was loaded at a flow rate of 0.8 mL/min and elution was monitored by UV absorbance at 280 nm.

### Cross-linking assay

100 *µ*L of Ox→ Ox and Red→Ox samples were prepared, with the denaturaiton step performed in 8M urea. Each sample were then split into two 50 *µ*L sub-samples, one of which was mixed with LDS loading dye + iodoacetamide (IAA) to a final concentration of 10 mM (Ox→ Ox and Red →Ox lanes)while the other was mixed with loading dye, IAA, and TCEP at a final concentration of 5 mM (Ox→ Ox reduced and Red→Ox reduced lanes). The samples were incubated in the dark for one hour to allow IAA to alkylate free cysteines (to prevent disulfide scrambling) and TCEP to reduce disulfides in the reduced sub-samples. All sub-samples were then loaded into an SDS-PAGE gel (∼5 *µg* protein per lane), which was run at 150V for one hour and fifteen minutes. The gel was then stained with Coomasie-G250 dye, photographed, and brightness and contrast were adjusted uniformly across the image to maximize intensity of bands.

### Atomistic simulations

Atomistic simulations were run using MCPU software [50] with enhanced sampling and replica exchange as descried previously [49]. MCPU uses a knowledge-based potential [73] to model (non-covalent) interactions between atoms, as well as torsional and bond-angle terms. In addition, in constructs with constraints, a harmonic term with respect to cysteine-cysteine distance, centered at a alpha-carbon distance of 5 Angstroms, was added for all native cysteine pairs. The initial structure of RBD was prepared from 334 to 528 of SARS-CoV-2 S1 protein (PDB ID: 6YOR), which was equilibrated for 50 million Monte-Carlo (MC) steps in our potential. The resulting energy-minimized structure was then used for a production run of 500 million MC steps, the last 400 million of which were used to compute equilibrium properties–at this point simulations were judged to be well converged.

To compute equilibrium probability distributions for a desired variable *X* (e.g., topological configuration, native contacts or inter-cysteine distance), the value of *X* was extracted for all simulation snapshots. In the case of topological configurations, substructures were created from the RBD contact map and snapshots were assigned to configurations as described previously [49]. For variable *X* of interest, we then computed a potential of mean force (PMF), or free energy as a function of *X* using the Multi-Bennett Acceptance Ratio (MBAR) method [74]. From these free energy profiles, the Bolztmann probability distribution over *X* can readily be computed and the mean of this distribution can be obtained. To ensure robustness, most quantities were computed at multiple simulation temperatures below the temperature at which the RBD was observed to melt.

Co-translational folding simulations were run and analyzed in much the same way as for full-length RBD, described above except for the design of the constraints. For constraints of co-translation folding simulations, the harmonic term was centered at a beta-carbon distance of 4.6 Angstroms, instead of alpha-carbon. The RBD was truncated at the end of the nearest secondary structure element following each cysteine of interest (Cys 3,28, 58, 99, 155, and 195) resulting in the chain lengths indicated in Fig. 5. We likewise ran simulations at each length, except the first, in which all prior cysteines except the most C-terminal one were constrained to their native disulfide pairs–these results are shown in SI.

### Kinetic model

To build a kinetic model, we define a set of states {*s*}, each of which corresponds to the collection of microstates in which some subset of observed disulfides are formed. A state is defined as containing either any permutation of the four native disulfides 3-28, 46-199, 147-155, and 58-152, or the observed nonnative disulfide 28-58 irrespective of whether other disulfides are present– this nonnative state is treated as an unproductive absorbing boundary. We assume state *s* can convert to state *s*^*′*^ if *s*^*′*^ contains all the same disulfides as *s* plus one additional disulfide *d*, and we assume that formation of *d*, and thus this *s* to *s*^*′*^ transition is irreversible. Although this is not strictly true given that disulfides can isomerize, the timescales for isomerization are relatively slow and may compete with nonnative aggregation–another absorbing boundary not considered here. For disulfide *d* to form starting from state *s* at translation length *L*, we require two things to happen, namely 1.) the cysteines comprising *d* need to be brought into contact (defined as a distance less than 5 Angstroms) via structural fluctuations, which occurs with equilibrium probability 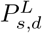 at length *L*, and 2.) The cysteine pair must be oxidized by PDI–a process that, for simplicity, is assumed to occur at a rate *k*_*ox*_ that is independent of length and disulfide pair identity. We also make the additional assumption that, at a given length, the probability of any cysteine pair forming is independent of whether other disulfides have already formed, and thus we can write 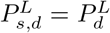 –this is justifiable because cysteine pairs in RBD are located in distinct structural regions. We further justify this assumption by running simulations up to each length at which a new cysteine pair has been synthesized in which all previous cysteine pairs have been constrained to form disulfides, and we observe that these constraints do not significantly affect the probability that the new pair forms (SI). We also assume that structural fluctuations are rapid compared to disulfide oxidation such that the effective disulfide-formation rate can be modulated by the equilibrium constant involving the probabilities that the cysteines are paired–this is akin to EX2 kinetics in hydrogen-exchange experiments.

Under these assumptions, the effective formation rate for native disulfide *d* at length *L* is given by equation (1). Likewise, all states in which neither disulfides 3-28 nor 58-192 have formed are assumed to be capable of transitioning to the nonnative state, indexed by *n*, in which the the 28-58 disulfide at a rate given by

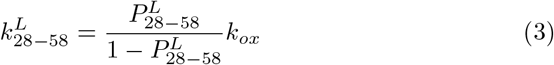

We then construct a transition matrix **M**^**L**^ corresponding to translation length L with entries given by:

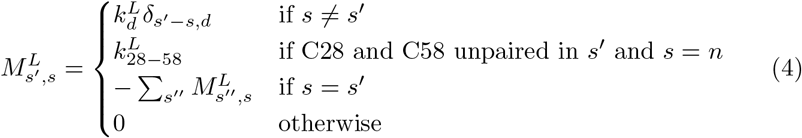

Where *δ*_*s*_*′−s,d* is a Kronecker delta function that takes on value 1 only if *s*^*′*^ contains all the same disulfides as *s* plus additional disulfide *d*. We note that this transition matrix conserves probability, and thus all columns sum to 0.

We let *P*^*L*^(*t*) denote the vector of probabilities of occupying different states at time *t* at translation length *L*. This vector satisfies the master equation(equation (2)):

To predict the time evolution of the disulfides while accounting for changing translation length, we first solve equation (2) numerically at a given translation length L, using the cysteine-pairing probabilites drawn from the simulation at that length, for an amount of time given by *τ*_*L*_ = *τ*_*AA*_* (*L*^*′*^− *L*), where *τ*_*AA*_ is the average time to translate 1 amino acid, and *L*^*′*^ is the next translation length simulated. The resulting probabilities, *P*^*L*^(*τ*_*L*_), are then used as the initial probabilities at the next length *L*^*′*^. At the first length *L* = 37, the initial probability is assumed to be 1 at the state with no disfulides and 0 at all other states. Once the final length is reached, the time evolution is solved for an additional amount of time shown in the figures.

## Acknowledgments

The authors thank David Thorn and Sourav Chowdury for technical assistance with experiments and useful discussions. The study was supported by the NIH-5R35GM139571-02. All computations in this work were run on the Harvard Canon Cluster. AB acknowledges funding from the National Science Foundation Graduate Research Fellowship Program (DGE1745303), The NSF-Simons Center for Mathematical and Statistical Analysis of Biology at Harvard (Award Number #1764269), and the Harvard Quantitative Biology Initiative.

## Authors’ contributions

A.B., K.P., and E.S. designed research. A.B. and K.P. performed research and analyzed data. A.B., K.P. E.S., and E.I.S. wrote the manuscript.

