## Supplementary Information for "Cotranslational formation of disulfides guides folding of the SARS COV-2 receptor binding domain"

### Mechanisms underlying folding and disulfide formation in the SARS CoV-2 receptor binding domain

Supporting Information

### Extended Methods

#### Pegylation assay

240 $\mu$L aliquots of Ox$\rightarrow$Ox and Red$\rightarrow$Ox refolded RBD samples at 2 $\mu$M were prepared, with denaturation performed using 8M urea, as well as an equivalent sample that had been treated with 1 mM TCEP during both denaturation and refolding steps (corresponding to "reduced" gel lane). Refolding was quenched after 6 hours via addition of urea to a final concentration of 7.6M (+1 mM TCEP in the case of "reduced" sample) and all samples were then concentrated down to 28 $\mu$L using Amicon Ultra 0.5 mL centrifugal filters. To each sample, 4 $\mu$L of mPEG-Maleimide (Creative Pegworks, average MW of 5 kD) dissolved in DMSO was added to a final concentration of 5 mM. An unpegylated control sample consisting of 28 $\mu$L native RBD (9 $\mu$M in 6M urea) and 4 $\mu$L of DMSO (no PEG-Mal) was also prepared. All samples were incubated for 2 hours at 37C, then the PEG-Mal reagent was removed using 2 mL Zeba spin desalting columns. 10 mM DTT was then added to fully reduce all samples. Samples were then mixed with NuPAGE LDS sample buffer to a final concentration of 1X, fully denatured at 95C for 15 minutes, then loaded onto a 4-12 % Criterion-XT gel (45 $\mu$L well volume). The gel was run for 60 minutes at 150V, and stained with Coomassie-G250 dye. The gel was then photographed, and brightness and contrast were adjusted uniformly across the image to maximize intensity of bands. Band intensity was then quantified using ImageJ software. To account for background and baseline effects, for every intensity peak, a straight line was manually drawn between the two intensity minima flanking the peak, and the area of the shape enclosed by this line and the intensity profile was measured. All areas for peaks corresponding to a given sample were then normalized by the total to obtain the fractional intensities shown in Fig. 3C. Peaks were only included if their height was visibly deemed to be above background fluctuations. Intensity profiles are shown in Supplementary Figure 5.

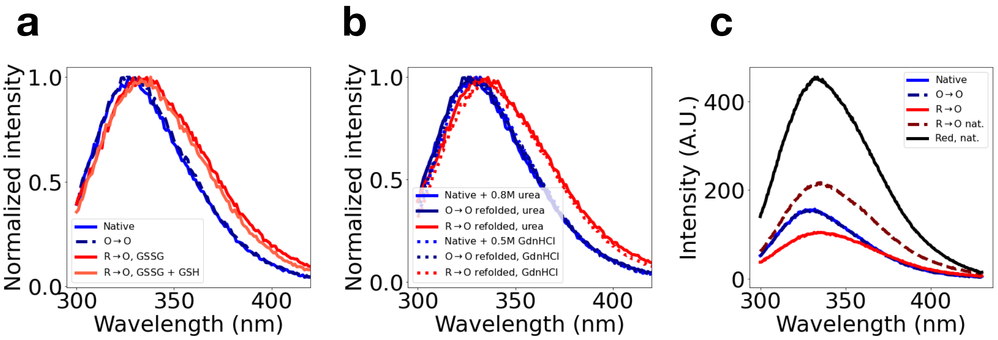


**Supplementary Fig. 1: Nonnative Red→Ox RBD spectrum is robust to gluathione buffer composition and denaturant used.** **a** Normalized fluorescence spectrum for native and Ox→Ox RBD, as well as Red→Ox RBD refolded in 5 mM oxidized glutathione (GSSG) buffer (as in main text Fig. 1) and buffer composed of 1.3 mM reduced glutahtione (GSH) + 2.7 mM GSSG. **b** Normalized fluorescence spectra of refolded Ox→Ox and Red→Ox RBD spectra, as in main text Fig. 1, in which denaturation step is performed in either 8M urea or 5M GdnHCl, as indicated in legend. In addition, native RBD spectra are shown in the presence of either 0.8M urea or 0.5M GdnHCl to match the final denaturation concentration in refolded samples. **c** Same as main text. Fig. 1c except spectra are not normalized.

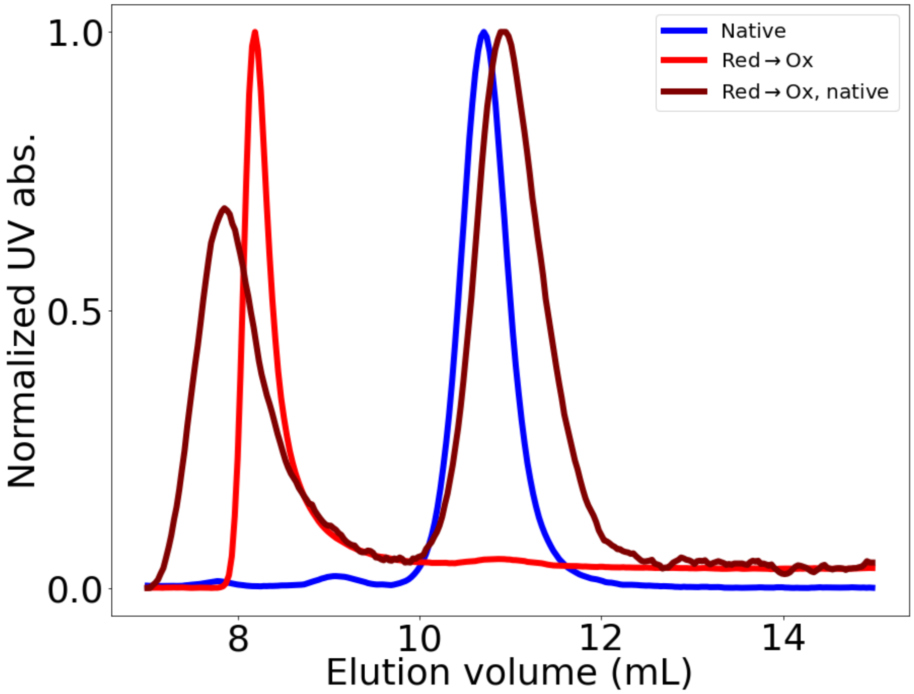


**Supplementary Fig. 2: Red→Ox preparation produces large, soluble aggregates regardless of whether reduction step is performed under native or denaturating conditions.**. SEC traces, obtained as in Fig 2 in the main text, are shown for native, Red→Ox, and Red→Ox native constructs. All constructs were prepared as described in the main text to a protein concentration of 10-20 $\mu M$. The weaker void volume (aggregate) peak and higher monomer peak observeed in the Red →Ox native trace may be a result of incomplete disulfide reduction under native conditions .

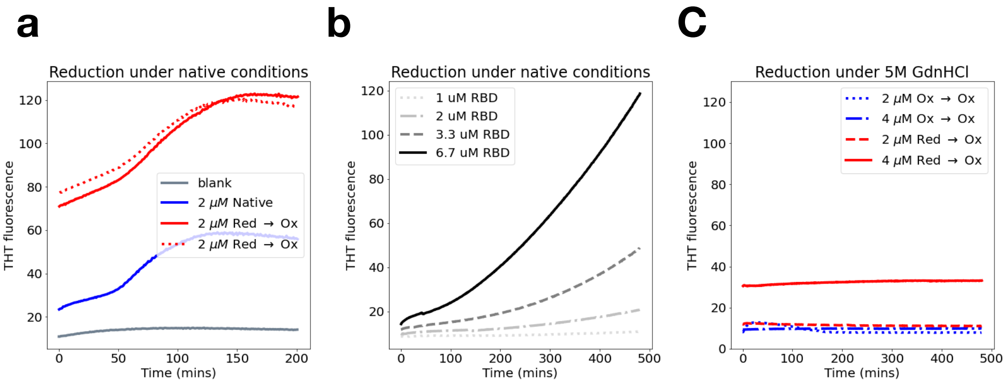


**Supplementary Fig. 3: Red→Ox construct shows heightened THT fluorescence relative to Ox→Ox under native conditions. a.** THT fluorescence traces as a function of time at 37°C for native RBD, two Red→Ox replicates in which reduction was performed under native (non-denaturing) conditions (see main text), and glutathione buffer blank. In these experiments $t = 0$ corresponds to the time at which THT was added and the temperature was increased to 37°C; reduction and re-oxidation were performed prior to the start of the experiment. **b.** THT fluorescence traces as a function of time at 37°C for RBD sample that was reduced under native conditions via addition of 1 mM TCEP at time 0. Protein concentrations are indicated in legend **c.** THT fluorescence traces as a function of time at 37°C for Ox→Ox and Red→Ox constructs in which reduction was performed under *denaturing* conditions. Protein concentrations are indicated in legend. The initial time, $t = 0$, is defined as in a., and reduction/re-oxidation were likewise performed prior to the start of the experiment.

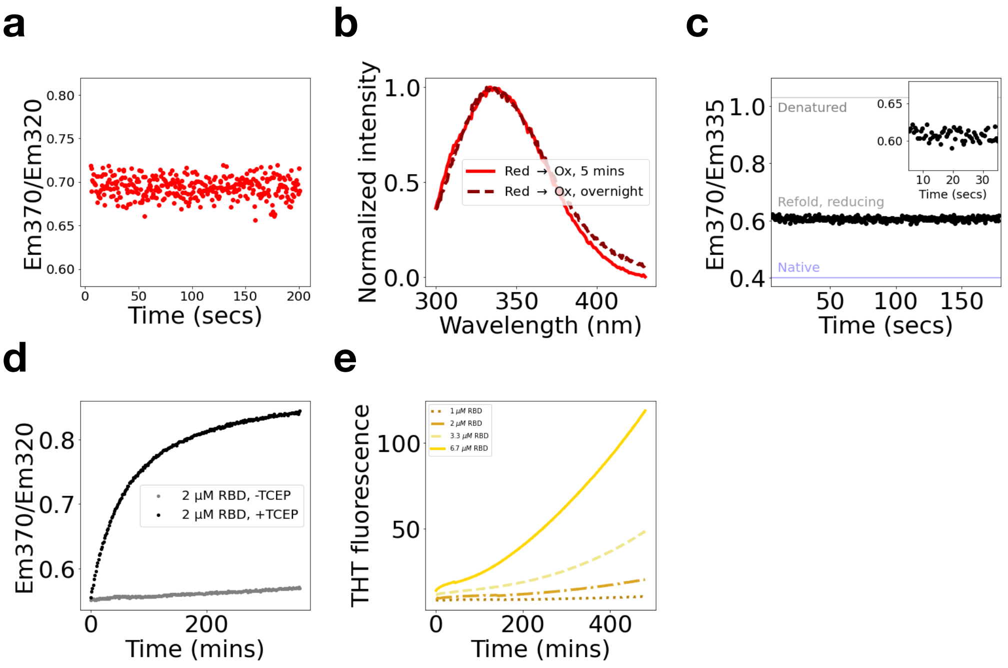


**Supplementary Fig. 4: RBD nonnative refolding is much faster than aggregation. a** Same as main text Fig. 2d, except we now report on the ratio of fluorescence intensity for an alternative pair of wavelengths–370 nm and 320 nm, rather than 370 nm and 335 nm, as in the main text. **b.** Normalized Red→Ox fluorescence spectra after 5 minutes of refolding (red) and after overnight refolding (brown) nearly overlap, suggesting kinetics are mostly complete after 5 minutes. The overnight sample shows somewhat higher fluorescence at wavelengths above 370 nm, which may be due to aggregation occuring on a slower timescale. **c** Same as main text Fig. 2d, except that refolding is now performed under *reducing* conditions (1 mM TCEP). We still observe a nonnative final ratio, consistent with main text Fig. 1, as well as kinetics that are too rapid to correspond to aggregation as argued in the main text, strongly indicating that a nonnative monomer fold is thermodynamically favored under these reduced conditions (Scenario 2 in main text). (d) Ratio of fluorescence emission at 370 nm to that at 320 nm as a function of time following initiation of reduction with 1 mM TCEP (black curve labeled ”+ TCEP”) or a control in which an equivalent volume of water is added (gray curve labeled ”+ TCEP”). The kinetics are slower than in panels a and c as nonnative misfolding is now rate-limited by reduction. (e) ThT fluorescence as a function of time (same as Supplementary Fig. 3b) following initiation of reduction as in panel d, at protein concentrations indicated. We observe that the timescales for ThT fluorescence increase–likely reflecting aggregation due to the pronounced concentration dependence–are much slower than the nonnative misfolding timescales, consistent with our interpretation of results shown in main text Fig. 2d and panels a and c above–namely that the nonnative fluorecence ratio that rapidly emerges after refolding corresponds to nonnative *monomer* folding, and aggregation happens more slowly thereafter.

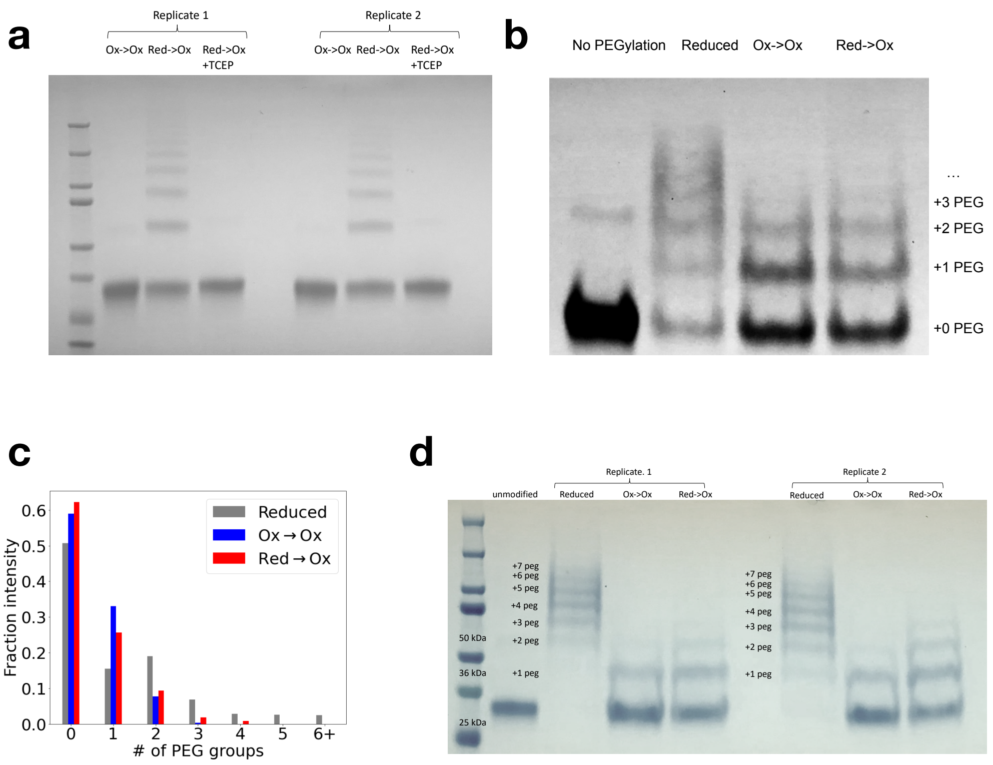


**Supplementary Fig. 5: Re-oxidized protein shows incomplete disulfide formation whether reduction step is performed under denaturing or native conditions**. **a.** Two experimental replicates of cross-linking gel assay, as per main text Fig. 3b, were performed. But unlike in the main text, the reduction step was now performed under native conditions(as per the Red→Ox, native and Reduced native protocol in Fig. 1). Both experimental replicates were loaded on an SDS-PAGE gel and stained with coomassie G-250 dye as in Fig. 3. Each lane is labeled with its corresponding sample. **b.** A PEGylation assay was used to assess differences in cysteine accessibility between different constructs–for details see extended method. PEGylated samples were loaded on an SDS-PAGE gel which was stained with coomase G-250 dye and shown here. Lanes correspond to (left to right) unpegylated RBD, fully reduced, Ox→Ox, and Red→Ox pegylated RBD samples. Bands corresponding to different numbers of PEGylations are labled on the Red→Ox lane. D**c.** Quantification of band intensity from SDS-PAGE gel in b. using ImageJ for Ox→Ox, and Red→Ox samples, normalized so that intensities for a given sample sum to 1. We note that bands with 6 and more pegylations overlap on the gel and thus cannot be resolved **d.** Two experimental replicates of the pegylation assay in panel b., were performed, but with reduction performed under native conditions (as per the Red→Ox, native and Reduced native protocol in Fig. 1). The gel was loaded and stained as in panel a, and all lanes are labeled with corresponding samples.

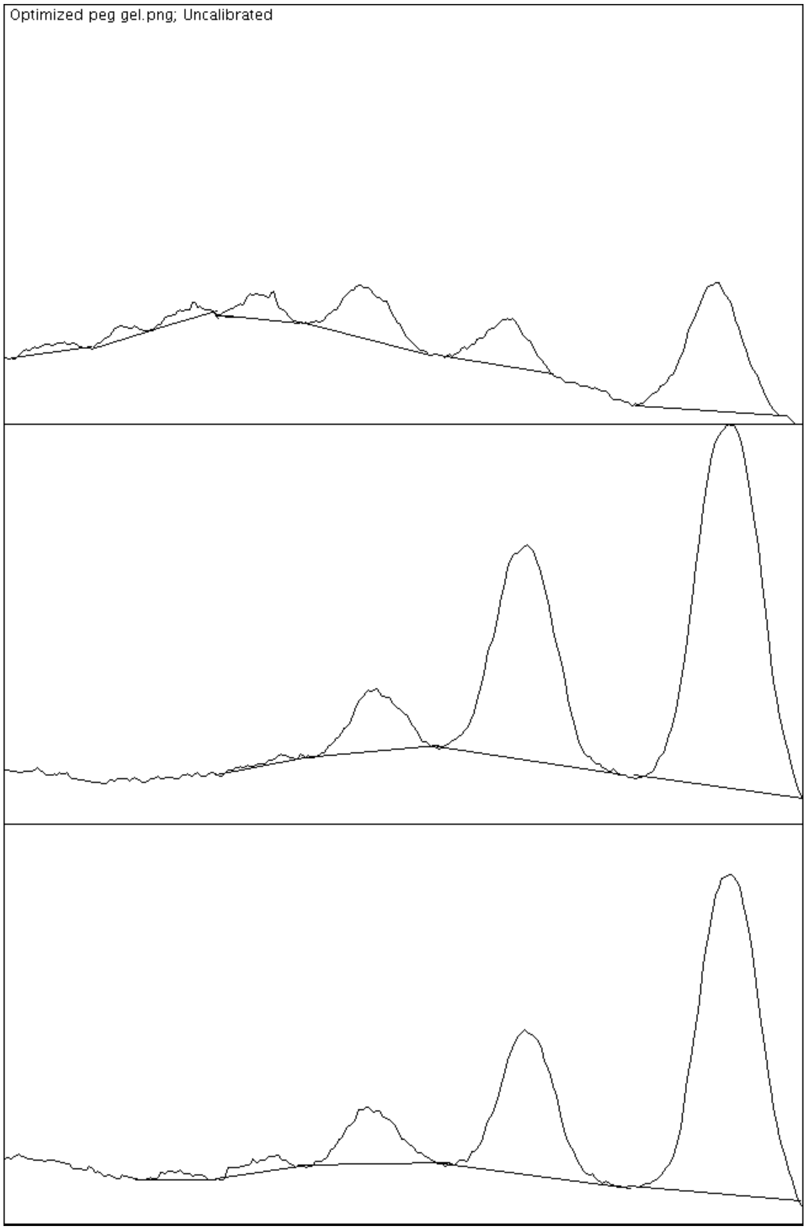


**Supplementary Fig. 6: Intensity plots for PEGylation gel.** Profiles of band intensity from PEGylation gel in previous figure, obtained from ImageJ software, are shown for Reduced (top), Ox→Ox (middle), and Red→Ox lanes. Baselines were drawn manually connecting adjacent minima, and the area between each baseline and peak was integrated. For details see extended methods.

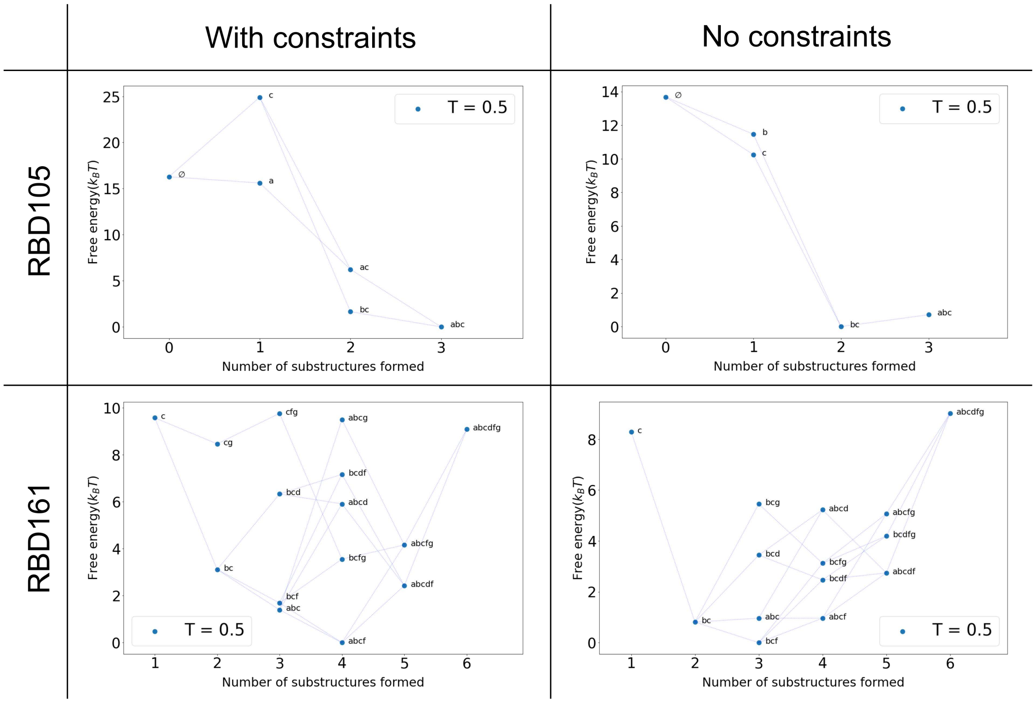


**Supplementary Fig. 7: PMF plots for intermeditate chain lengths.** Free energy difference (ΔG) with respect to the lowest free energy state as a function of a number of substructures formed for intermediate chain lengths (RBD105 and RBD161) with and without constraints. RBD105 has a fully-folded state as energy minimum in constrained simulation, while a partially-folded structure is slightly favored in non-constrained simulation. In contrast, for RBD161, partially-folded structures (substructure *abcf* for the simulation with constraint and substructure *bcf* for the simulation without constraints) are favored regardless of constraints.

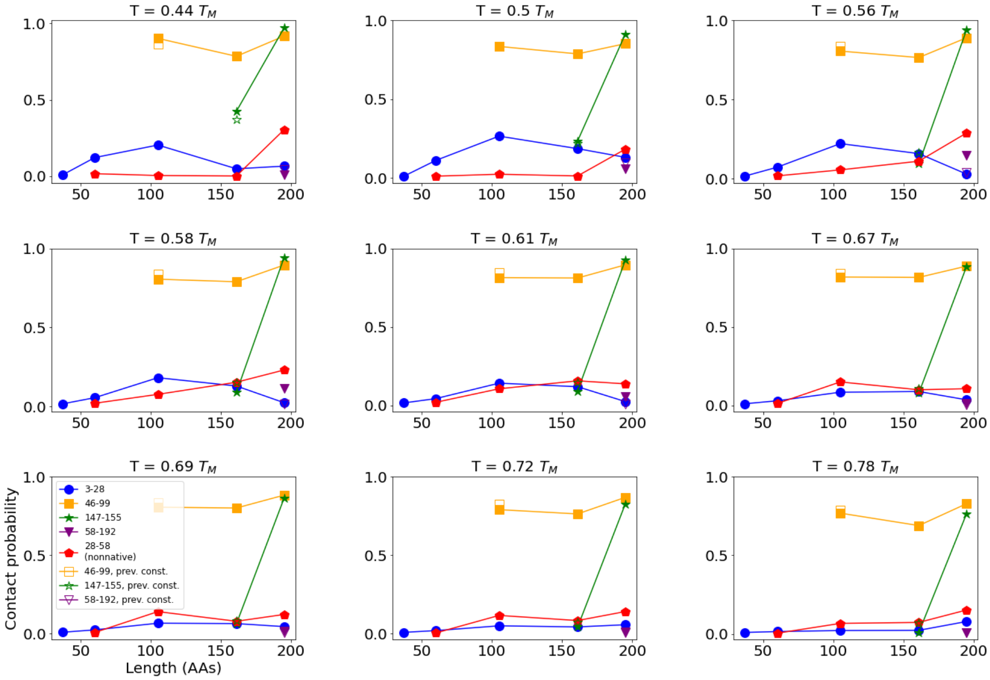


**Supplementary Fig. 8: Dependence of cysteine pairing probability on simulation temperature and existing disulfides.** These panels show the probability that the indicated cysteine pairs come within 5 Angstroms of each other, as per main text Fig. 5a, at various simulation temperatures indicated above the respective plots. In the context of our coarse-grained potential function, we cannot trivially map simulation temperature to real temperature–thus all temperatures are expressed in units of the melting temperature $T_M$. Although real physiological temperatures typically correspond to $\sim 0.95 T_M$ for most proteins, this is not necessarily the case in our potential function, which is temperature independent. Most panels in the main text are shown at $T = 0.56\ T_M$–a temperature at which the native state is predicted to be stable in the presence of constraints, and at which native cysteine pairs typically show high probability of coming together co-translationally. In addition to solid symbols, these panels also show cysteine pairing results from a control simulation in which, at the first length at which a native cysteine pair can form $d$, all previous native cysteine pairs are subject to disulfide constraints (hollow symbols, labeled "prev. const". ) For example, the points labeled "147-155", prev. const. (hollow green stars) give the probability that cysteines 147 and 155 make contact within a simulation at length 161 in which cysteines pairs 3-28 and 46-99 have been constrained to form disulfides. These results indicate that the presence of existing disulfides does not significantly change the probability that additional disulfides form, justifying a key assumption in our kinetic model (see main text).

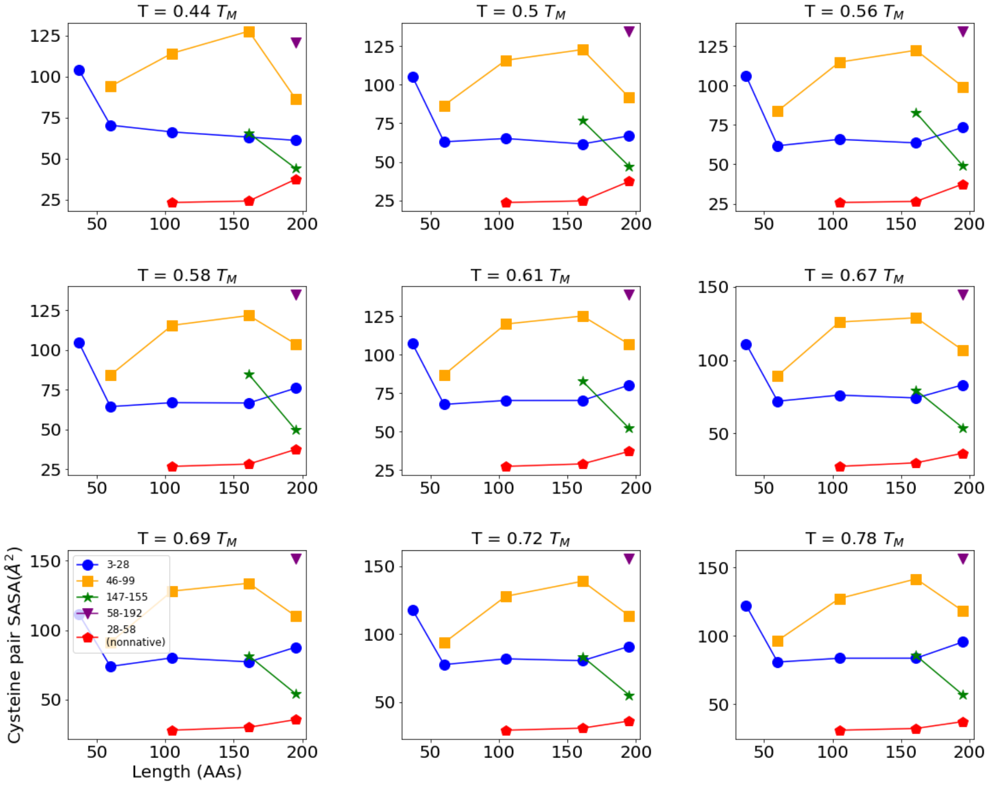


**Supplementary Fig. 9: Solvent-exposed surface area for cysteine pairs.** These panels show the total solvent exposed surface area (SASA) for different cysteine pairs, conditioned on those pairs being in contact, at different simulation temperatures (defined as per previous panel). In general, for a given pair, we observe the SASA shows minimal depenence on length, particularly at lengths where the pairing probability (shown in previous figure) is appreciable.

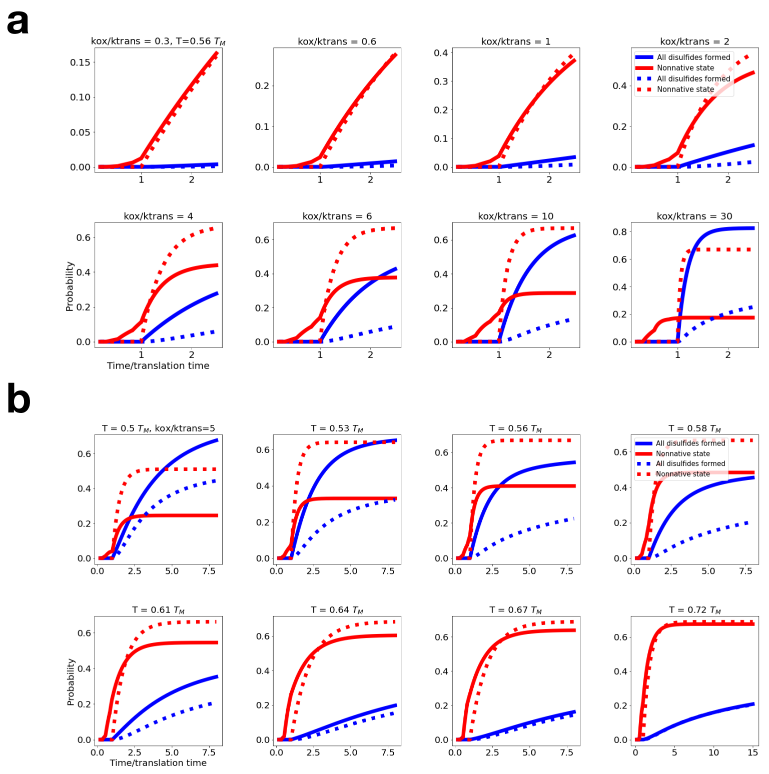


**Supplementary Fig. 10: Dependence of kinetic model results on rate and temperature parameters.** **a.** As in main text Fig. 5, the probability, as a function of time normalized by translation time, of occupying the state in which all disulfides are formed (blue) and that in which the nonnative 28-58 disulfide is present (red) is shown for different values of $k_{\rm ox}/k_{\rm trans}$ indicated above the respective plots, all at a temperature of $T = 0.56\ T_M$ (temperature defined as in previous figure). As in the main text, we either allow for the possibility of co-translational disulfide formation (solid lines) or assume disulfide formation does not begin until the domain is fully synthesized (dashed lines). Co-translational folding is most advantageous for large values of $k_{\rm ox}/k_{\rm trans}$ as the nascent chain can optimally exploit intermediate lengths at which the $3-28$ disulfide is more likely to form. However, the largest value shown ($k_{\rm ox}/k_{\rm trans} = 30$ is unlikely to be physiologically reasonable as, assuming a $\sim 20\%$ co-translational cysteine pairing probability, the effective oxidation timescale would be 6 times the translation timescale, (i.e. about seconds ). This is much faster than measured PDI enzymatic timescales. **b.** As in main text Fig. 5, the probability, as a function of time normalized by translation time, of occupying the state in which all disulfides are formed (blue) and that in which the nonnative 28-58 disulfide is present (red) is shown assuming diffetent simulation temperatures, assuming a value of $k_{\rm ox}/k_{\rm trans} = 5$. In main text Fig. 5, a temperature of $T = 0.58\ T_M$ is used. The results are somewhat sensitive to temperature, as cysteine pairing probabilities tend to be higher at lower simulation temperatures, cysteines are more likely to pair up (see previous figure). At higher temperatures, co-translational folding still slightly improves the native folding yield by decreasing the nonnative fraction containing the $28-58$ disulfide, but this effect mostly vanishes by $T = 0.72\ T_M$
